# Structural Characteristics and Proton Conductivity of the Gel Within the Electrosensory Organs of Cartilaginous Fishes

**DOI:** 10.1101/2021.01.04.425255

**Authors:** Molly Phillips, Alauna Wheeler, Matthew J. Robinson, Valerie Leppert, Manping Jia, Marco Rolandi, Linda S. Hirst, Chris T. Amemiya

**Affiliations:** Department of Biology, University of Washington, Seattle, WA 98195, USA; Department of Molecular and Cell Biology, University of California, Merced, Merced, CA 95343, USA; Department of Physics, University of California, Merced, Merced, CA 95343, USA; Department of Materials Science and Engineering, University of California, Merced, Merced, CA 95343, USA; Department of Electrical and Computer Engineering, Baskin School of Engineering, University of California, Santa Cruz, Santa Cruz, CA 95064, USA; Quantitative and Systems Biology Program, University of California, Merced, Merced, CA 95343, USA

## Abstract

The Ampullae of Lorenzini (AoL) of cartilaginous fishes are sensory organs used to detect environmental electric fields. The proximal ends of the organs are externally visible as pores in the skin that lead into gel-filled tubular canals which terminate in rounded chambers filled with highly specialized electrosensory cells. The viscoelastic gel that fills the organs is composed of proteins and polysaccharides that are not yet completely characterized but are thought to play a critical role in the electrosensing mechanism. Although recent studies have identified various components of AoL gel, it has remained unclear how the component molecules are structurally arranged and how their structure influences the overall function of the AoL. Here we present the first microscopic descriptions and x-ray scattering data from AoL gel extracted from spotted ratfish (*Hydrolagus colliei*). Our results suggest that AoL gel is colloidal in nature and composed of spherical globules that are approximately 10-100 nm in size. We investigated the structural influence of the protein components of the gel specifically by analyzing gel that had been digested *in situ* via enzymatic proteolysis. By comparing gel before and after digestion using microscopy, x-ray scattering analyses, and proton conductivity measurements, we directly observed the structural and functional influence of proteins in AoL gel. The findings described here represent the first detailed structural analysis of AoL gel and lay the groundwork for more detailed studies into the specific interactions of molecules inside AoL gel at the nanoscale, with particular reference to their mechanistic role in electrosensing.

## Introduction

Electroreception, the ability of some animals to detect electric fields, is widespread amongst vertebrates. Some of the most well-studied electroreceptive animals are rays, skates, sharks, and chimaeras – cartilaginous fishes of the class Chondrichthyes. These fishes use specialized electrosensory organs for the detection of low frequency electric fields from biological sources such as prey or mates, and even for navigating using earth’s geomagnetic field (1-3). These electrosensory organs, called Ampullae of Lorenzini (AoL), are observable externally as small pores that are open to the surrounding environment (Figure 1A). In sharks and chimaeras, pores are most concentrated on the snout and around the mouth whereas in skates and rays, they are more widespread across the ventral and dorsal surfaces of the animals (4, 5). AoL pores lead into tubular collagen-wrapped canals of varying length and diameter (depending on species and location on body) that are lined on the inside by two layers of epithelial cells. At the distal end of each canal is a bulbous alveolus containing electrosensory hair cells that synapse with neurons connected to the medulla (Figure 1B). Importantly, the organs are filled with a viscoelastic gel that can be extracted by applying pressure to the skin adjacent to AoL pores and subjected to biophysical examination (Supplemental Figure 1). While the sensory capabilities of the AoL have been studied in detail over the past several decades, the process by which signals move from the environment through the organs, and how the gel is involved, remains a subject of debate (3).

**Figure 1.**
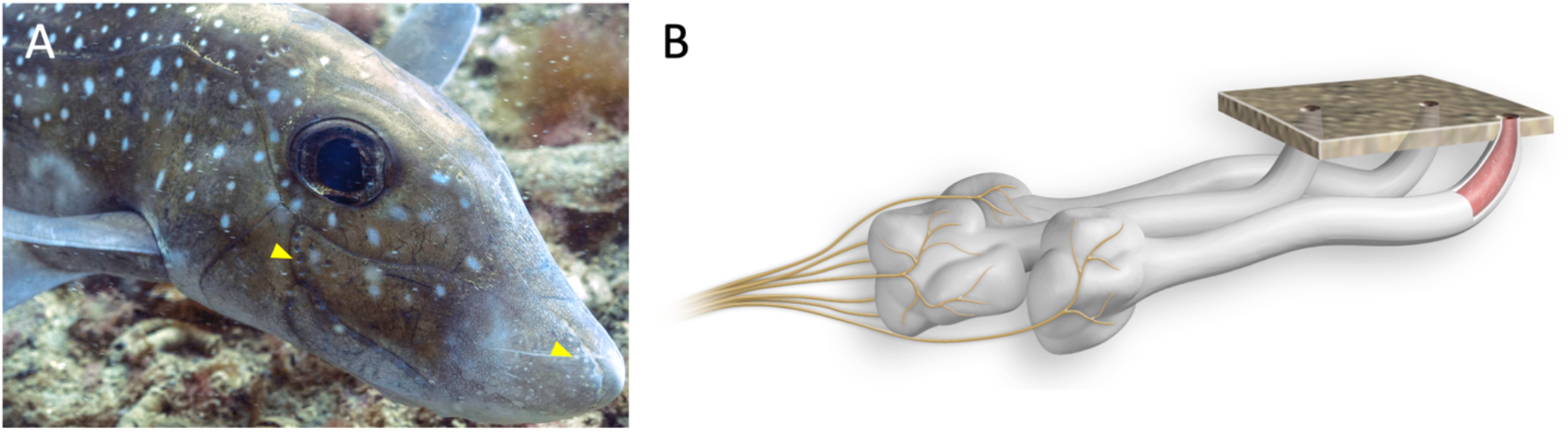
Anatomy of Ampullae of Lorenzini (AoL). **A)** An image showing a type of chimaera called a spotted ratfish (*Hydrolagus colliei*). Yellow arrowheads delineate the locations of some AoL pores. Image taken by Mick Otten. **B)** A diagram showing three AoL below the skin surface. Pores lead into canals filled with a gel (as made visible by the cutaway). Neurons (yellow) synapse with specialized electrosensory cells in the alveoli and project onto the hindbrain.

Research in the 1960s showed that AoL gel is approximately 95% water containing Na^+^, K^+^, Cl^-^, and urea (6). By measuring the relative abundances of hexosamines, sulphates, and other chemical groups, researchers identified that the gel contains various “mucopolysaccharides,” but their identities have remained unknown until recent years (7). AoL gel extracted from three fish species demonstrated extremely high proton conductivity, higher than that of any other reported biological material (8). It has been proposed that keratan sulfate, a glycosaminoglycan, is a component of AoL gel and responsible for conferring the substance’s proton conductive properties (8, 9). Various studies have attempted to uncover the complete molecular makeup of AoL gel, most with a focus on polysaccharides and proteins (9-11). Cellular contamination during AoL gel extraction, however, makes the interpretation of proteomic datasets challenging. We recently reported evidence suggesting that chitin is another polysaccharide component of the gel, but it remains unknown if the chitin molecules are modified in some way or complexed with other components in order to promote solubility in the aqueous gel (12).

Major gel-forming components have been shown to exist as fibrous or rod-like particles, flexible coils, spherical particles, or linear molecules held together by crystalline junctions (13). However, it is still unclear how the molecular components of AoL gel are arranged and how they contribute to the gelatinous nature of the material. Further, it is unknown how the gel’s viscoelastic structure factors into the proton conductivity of the substance. Using data from proteomics and polysaccharide analyses, Zhang *et al*. proposed a hypothetical model of AoL gel structure consisting of actin filaments holding together a scaffold of mucins bound to keratan sulfate molecules (9). Zhang *et al*. based their model on inferred interactions of gel components that they identified using chemical extraction methods and proteomics; the model was not based on biophysical structural data. Here, we describe an entirely different approach to studying the structure of AoL gel samples mostly from spotted ratfish (*Hydrolagus colliei*) (Figure 1A) using small angle x-ray scattering (SAXS), scanning electron microscopy (SEM), and atomic force microscopy (AFM). To understand the influence of the gel’s protein component specifically, we digested gel with the proteolytic enzyme, proteinase K, and compared the gross morphology and scattering properties of gel before and after protein removal. Further, we studied the influence of proteins on the proton conductivity of the material using electrochemical impedance spectroscopy. Our results, *in toto*, suggest a different model for the gel structure than had previously been suggested.

## Results

Low magnification fluorescence microscopy was used to study the structure of formalin-fixed tissues from little skate (*Leucoraja erinacea*) and spotted ratfish (*H. colliei*) that were stained with fluorescent probes consisting of the chitin binding domain (CBD) (12, 14) (Supplemental Figure 2A, B). CBD detects chitin within AoL gel material in a highly specific manner (12), and served as a marker of the fine outlines of the gel. While gel inside fixed whole-mount AoL exhibited wavy fabric-like patterns (Supplemental Figure 2A), the gel of sectioned AoL appeared to be organized into small packets that looked to be extruded from individual cells (Supplemental Figure 2B). Importantly, it is unclear how much of the observed structure resulted from fixation. To overcome this, we also stained native AoL gel extracted from *H. colliei* with CBD and imaged with fluorescence microscopy to reveal a sheet-like macroscopic structure (Supplemental Figure 2C, D). The shape of the gel’s structure was reminiscent of the wrinkled appearance of other biopolymer gels (15). All experiments described hereafter used native, unfixed gel material.

To study the structure of AoL gel at higher magnifications, we used AFM and SEM. When imaging *H. colliei* AoL gel with AFM, abundant spherical globules were observed at various locations across the mica surface (Figure 2A, B). These globular structures were often piled on top of one another (inset of Figure 2A) and in some cases, appeared to combine to form large aggregates. An average globule had a diameter of approximately 100 nm. The globules appeared to be closely associated and buried in a mat that covered the mica. When smaller AFM scan areas were used, rope-like objects could also be seen weaving in-between the globules (Figure 2B).

**Figure 2.**
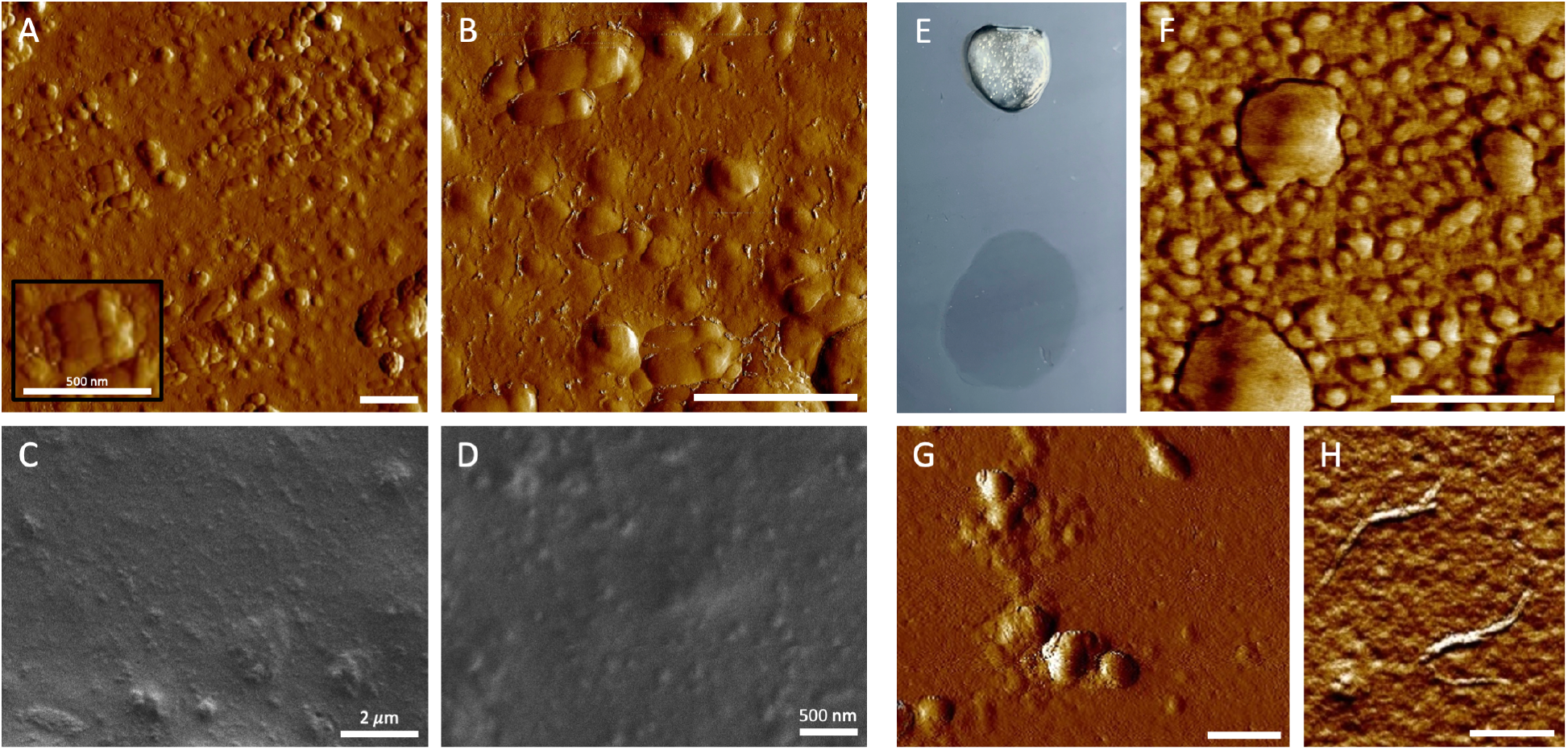
Microscopic analysis of AoL gel before and after digestion with proteinase K. **A, B)** AoL gel from *H. colliei* was diluted 1/20 in water, dropped on a freshly cleaved mica sheet, air dried and imaged with AFM. Globular structures were observed widespread across the surface (A) and close examination revealed rope-like objects winding in-between the globules (B). Inset in (A) shows organized packing of globules. **C, D)** Pellet resulting from ultracentrifugation of AoL gel was dragged across a freshly cleaved piece of mica then imaged with SEM. Images show same sample (in slightly different fields) at lower (C) and higher (D) magnification. The majority of the surface was covered in a mat of material, which at higher magnification, appeared to be composed of globules comparable in size to those observed with AFM. **E)** Image showing a fresh aliquot of *H. colliei* AoL gel (top) above a drop of proteinase K-treated *H. colliei* gel (bottom) demonstrating the reduction in material stiffness following protein digestion. **F)** AFM image showing a sample of *H. colliei* gel that had been digested with proteinase K, diluted 1/20 in water and dropped onto a freshly cleaved mica sheet to ambiently dry. We observed globules spread across the mica surface and like those globules observed in the non-digested gel samples. **G)** AFM image showing a sample of *H. colliei* gel that had been digested with proteinase K, dialyzed with 12-14 KDa tubing, then dropped and ambiently dried on freshly cleaved mica. Despite the definite reduction in protein material as evidenced by SDS PAGE (Supplemental Figure 4), aggregating globules were still clearly observed. **H)** AFM image showing whisker-like crystalline structures resulting from sonication of the same sample of proteinase K-digested AoL gel that is shown in (G). Scale bars – A, B, D, F-G: 500 nm; C: 2 *μ*m.

When freshly extracted gel was deposited on an aluminum stub, no consistent features were observed with SEM. Therefore, knowing that AoL gel is mostly water (6), ultracentrifugation was used to concentrate gel components and separate them from water. Pelleted AoL gel material was smeared across a piece of mica, then imaged with SEM at various magnifications (Figure 2C, D). At lower magnification, dense aggregated material was observed in dispersed mounds (Figure 2C). When higher magnification was used, globules of similar shape and size to those observed with AFM were resolved (Figure 2D). The shape of the objects observed with both AFM and SEM led us to suspect that AoL gel is colloidal in nature and composed of spherical particles. However, desiccation undoubtedly had some impact on the AoL gel structure, as drying effects have been shown to reduce the natural porosity of gel materials (16, 17).

To overcome the inevitable effects of desiccation on the AoL gel structure, we used SEM to image gel samples that had been exposed to supercritical CO_2_. The supercritically dried sample exhibited a fine porous network similar in form to published images of so-called aerogels (Supplemental Figure 3A) (18, 19). Aerogels are produced when hydrogel is dried in a manner that allows for the maintenance of the original form, such as by supercritical drying (20). Given this, the SEM images of supercritically dried gel are likely more representative of aqueous gel structure than those SEM images of dried and vacuum dessicated gel. Close examination of the supercritically dried gel structure revealed various structural arrangements, some densely packed and solid (Supplemental Figure 3B), but most porous and loose (Supplemental Figure 3C). In all cases, the gel structure appeared to consist of globular elements that were on average 100-200 nm in diameter. These findings corroborated the proposition that AoL gel is colloidal and composed of aggregating spherical objects.

Next, we wanted to investigate which macromolecular gel components comprised the spherical globules observed by microscopy. Knowing that AoL gel is composed primarily of polysaccharides and proteins, we digested aliquots of gel from *H. colliei* with proteinase K and studied the properties of the material that remained after digestion. Remarkably, after one hour of proteinase K exposure, the AoL gel transitioned into a fluid state. This phenomenon was demonstrable when samples were dropped onto a glass slide before and after digestion (Figure 2E). To assess the efficacy of enzymatic digestion, we ran samples of native and digested AoL gel on polyacrylamide gels and observed a marked reduction in detectable protein before and after treatment with proteinase K (Supplemental Figure 4). Dialysis with an aliquot of digested AoL gel using ∼14 KDa MWCO tubing revealed an even greater reduction of proteins (Supplemental Figure 4). When digested gel was imaged with AFM, we observed globules similar to those that were seen comprising native AoL gel (Figure 2F, G). These globules were visible both in undialyzed (Figure 2F) and dialyzed (Figure 2G) samples. Additionally, when a sample of proteinase K-digested gel was sonicated prior to AFM imaging, the material took on an entirely new form (Figure 2H). Interestingly, the structures resulting from sonication resembled published images of chitin crystals generated by chemically extracting polysaccharides from AoL gel (Figure 2H), indicating that chitin likely composes at least some portion of the globules (12).

Microscopic evidence suggested that samples of native and proteinase K-digested AoL gel were composed of aggregating spherical particles. However, whereas globules appeared to be in tight mat-like association in native gel samples (Figure 2A, B), they looked more dissociated in digested gel (Figure 2F, G). Therefore, we next asked whether there were differences in the internal structure of the globules in aqueous solutions of native and digested AoL gel. To address this question, we performed SAXS on AoL gel before and after proteinase K digestion. SAXS analyses are ideal for studying poorly ordered dilute aqueous materials like AoL gel (21). As with the samples used for SEM, we first ultracentrifuged the gel to concentrate the major macromolecular components. Ultracentrifugation of native gel yielded a thick white pellet while digested gel formed a yellowish cloud at the bottom of the centrifuge tube. Both tubes contained a sizeable proportion of clear supernatant. The supernatant and the concentrated material from both samples were placed in a thin-walled x-ray capillary tube before exposure to the x-ray beam. Figures 3A and B show SAXS scattering patterns from native gel and proteinase K-digested gel respectively. The supernatant from both gel samples exhibited scattering patterns extremely similar to water, whereas the concentrated materials displayed some notable differences (Figures 3A, B). Differences were especially apparent after background scattering from water was subtracted from both samples. The resulting curves reflect scattering primarily from polymers and proteins existing in the gel materials (Figure 3C).

**Figure 3.**
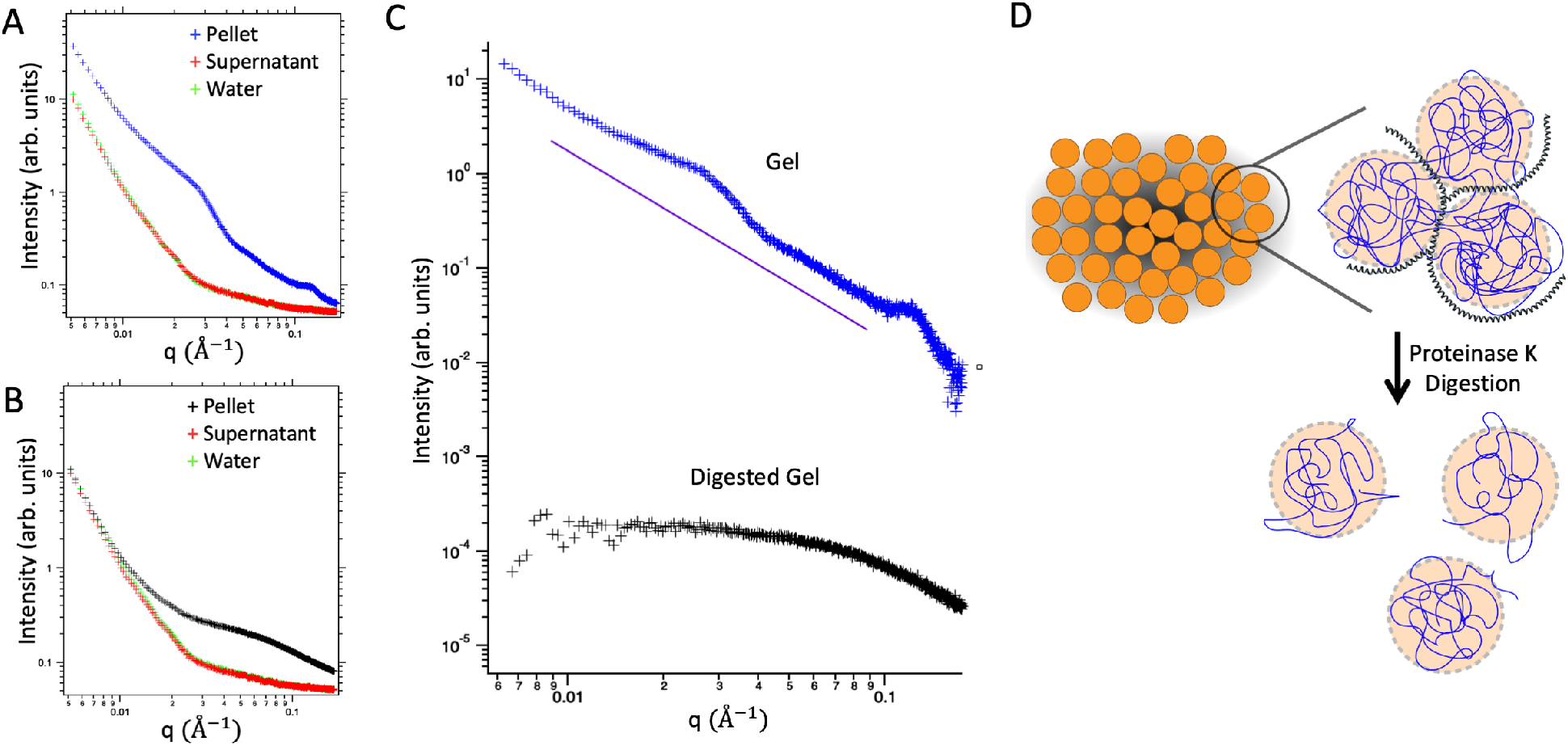
Comparison of native *H. colliei* AoL gel and proteinase K-digested AoL gel using SAXS. Scattering intensity is plotted as a function of the scattering vector, *q*. **A, B)** SAXS data using material resulting from ultracentrifugation of untreated AoL gel (A) and proteinase K digested AoL gel (B). **C)** Comparison of SAXS data from native gel pellet and the digested gel’s light cloudy material. Both plots are shown with water background subtracted. The purple line indicates a slope of −2 as a guide to the eye. **D)** Hypothetical model of AoL gel structure based on microscopy and SAXS. Globules observed with SEM and AFM are depicted in orange. Polysaccharide polymers (blue) are packed into rough spherical shapes that are held in close association by proteins (represented by black coils) when gel is in its native state (top). Digestion with proteinase K breaks up the majority of these proteins (bottom) which leads to fluidization of the gel.

The scattering curve resulting from undigested AoL gel has a slope close to −2 on a log-log plot (Figure 3C, blue plot). The magnitude of the slope suggests that AoL gel components are organized similarly to an ideal polymer chain over length-scales from ∼90 – 6 nm. An ideal (Gaussian) polymer chain is freely jointed and self-avoiding, with characteristic scattering described by the Debye function, S(q) μ 1/q^*v*^ where *v* is exactly equal to 2 (22). For a self-avoiding chain, the value of *v* is slightly higher as the chain effectively swells. The purple line in Figure 4C shows a slope of −2 as a guide to the eye. The curve has a clear broad peak at *q*= 0.028 Å^−1^, which corresponds to a real space lengthscale, *d* of 22 nm (where 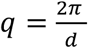). This peak may represent a lengthscale related to the average separation of the globules in the gel. A second peak can also be observed at approximately 0.1 Å^−1^, corresponding to a real space length-scale of *d*= 5 nm and we hypothesize that this peak is explained by the presence of some cellular debris within the gel, namely cell membranes, which are on average approximately 5 nm in thickness (23).

In contrast, the concentrated proteinase K-digested sample produced a scattering curve with no obvious peaks that levels off at a low *q*, indicating a lack of large-scale aggregated structures in the cloudy material resulting from ultracentrifugation. The curve is reflective of the material’s structure after digestion (Figure 3C, black plot) and suggests that the polymers remaining in the gel after digestion lack an organized arrangement and are likely dilute. A model summarizing our analysis of these findings is shown in Figure 3D and is consistent with our microscopy results.

Finally, in order to develop an understanding of the functional differences between AoL gel before and after proteolytic digestion, we compared the proton conductivity of native AoL gel from *H. colliei* and proteinase K-treated gel with alternating current (a.c.) electrochemical impedance spectroscopy (EIS) (Figure 4A). Both gel samples were dialyzed against H_2_O before measurements to remove other ions that could contribute to the conductivity. The Nyquist plots of both native and digested AoL gel show semicircles in the high frequency region and an inclined spur in the low frequency region (Figure 4B). These features are fingerprints of materials with predominant ionic conductivity (24). We calculated the conductivity using a simple equivalent circuit (Figure 4C) (25-27). For native AoL gel, we calculated the effective conductivity to be 5 ± 3.5 mS cm^-1^, which is consistent with the value of the conductivity (2 ± 1 mS cm^-1^) of AoL gel from other cartilaginous fish species measured previously (8) using transmission line measurement devices. Given the variability associated with samples extracted from biological specimens, the agreement of the values within the uncertainty range is remarkable. For the digested gel, we calculated the effective conductivity to be 22 ± 8.5 mS cm^-1^. The sizeable error observed with both measurements comes from the uncertainty associated in evaluating the thickness of gel material in the devices (Supplemental Figure 5). Regardless, it appears that the conductivity of the proteinase K-digested material is higher than that of native AoL gel. To further confirm that the charge carriers in the gel are protons, we studied the kinetic isotope effect by loading the gel with H^+^ and D^+^ in water (H_2_O) vapor and deuterium oxide (D_2_O) vapor, respectively (8). The resistance of both native and digested AoL gel in D_2_O vap0r was significantly higher than that in water as expected when protons dominate the conductance of the material, because the mobility of D^+^ is approximately half of the mobility of H^+^ (Figure 4D, E). This result provides confirmation that AoL gel from *H. colliei* is a proton conductor at high relative humidity and corroborates our previously reported observations (8).

**Figure 4.**
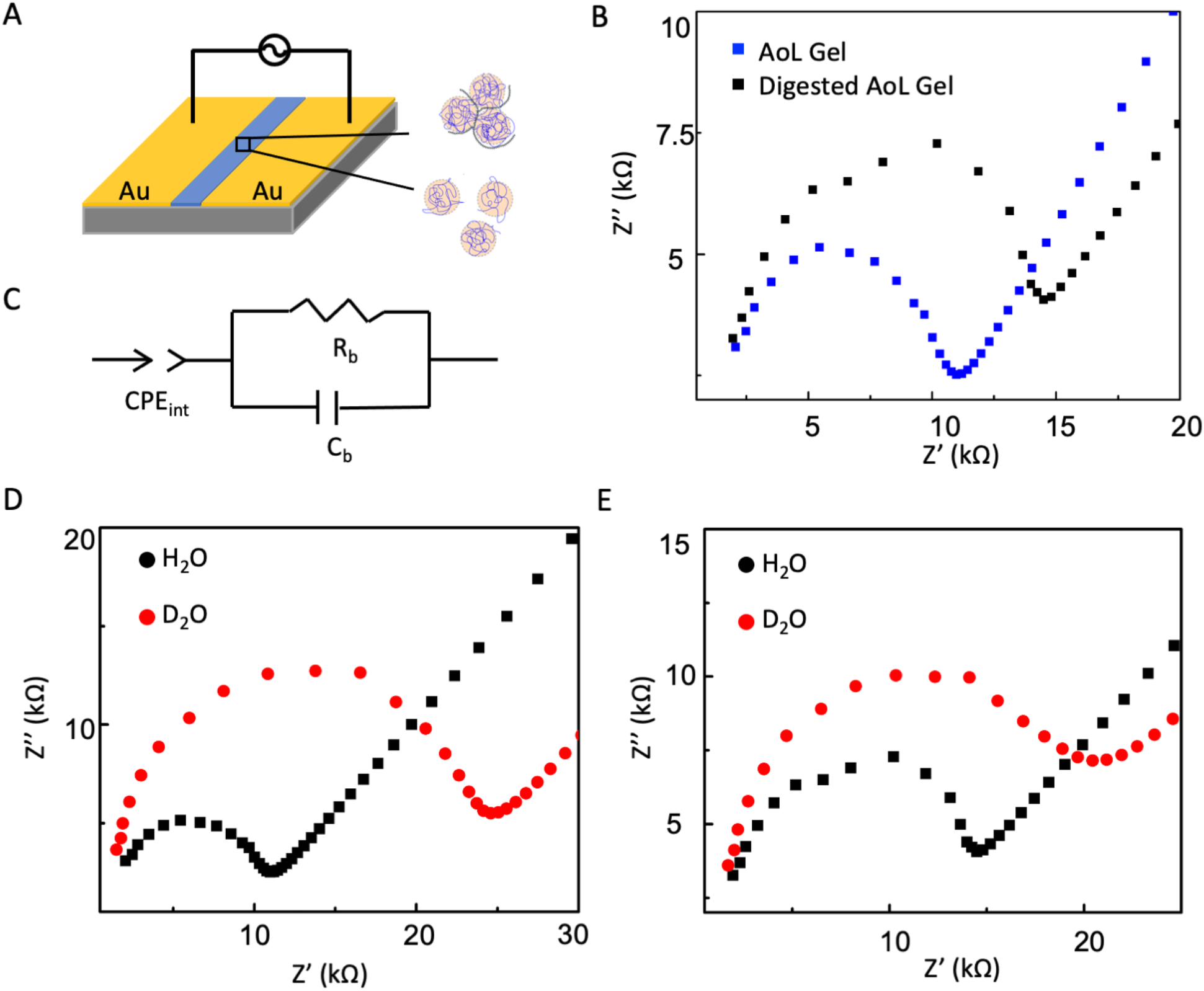
Proton conductivity of *H. colliei* AoL gel before and after proteinase K digestion. **A)** Two-terminal device used for EIS measurement. **B)** Nyquist plots of native AoL gel (blue) and digested AoL gel (black) at 90% relative humidity. **C)** Equivalent circuit model. A Constant Phase Element (CPE) was used to describe the non-ideal interface capacitance; Rb and Cb represent the resistance and capacitance of the sample, respectively. **D, E)** Nyquist plots of native *H. colliei* AoL gel (D) and proteinase K-digested *H. colliei* AoL gel (E) in the presence of H_2_O vapor (black) and D_2_O vapor (red).

## Discussion

Evidence from microscopic examination suggested that AoL gel from *H. colliei* is colloidal in nature, with spherical globules composing a substantial portion of the observable macromolecular structure. SAXS data using concentrated AoL gel revealed a scattering peak corresponding to a real space lengthscale of 22 nm. Globules were still observable by AFM in proteolyzed gel samples, whereas the 22 nm peak was completely absent from the SAXS plot generated using the same material. Taken together, these findings suggest that the 22 nm peak represents a lengthscale related to the average separation of the spherical globules within AoL gel. In addition, for the undigested samples, the range of lengthscales over which we observed close to ideal chain behavior (90 – 6 nm) suggests that there is some degree of chain interdigitation between globules in the aggregated structure, producing a continuous polymer network. It is important to point out that ultracentrifugation may have impacted the size of the polymer globules in native AoL gel therefore 22 nm would represent a lower correlation length limit.

When ambiently dried AoL gel was imaged with AFM, rope-like structures were observed weaving amongst the globules (Figure 2A, B). These structures were absent in proteinase K-digested gel, which suggests that they consist of proteins involved in holding the polymer network together. Elimination of these tethering proteins by enzymatic digestion seemingly collapsed the polymer network and although the globules remained, they became dispersed randomly in solution (diagrammed in Figure 3D).

It is still unclear what materials compose the globules entirely. The fact that the globules were still clearly resolvable after proteinase K digestion, makes it unlikely that they are highly composed of proteins. Therefore, the material remaining after proteinase K digestion could be composed of polysaccharides, lipids, or a combination of macromolecules. However, the globules comprising AoL gel resemble published images of chitosan nanoparticles (28); moreover, crystals reminiscent of chitin nanowhiskers were formed when the carbohydrate-extracted gel material was sonicated (29). These observations likely indicate that chitin makes up a substantial portion of the spherical globules, perhaps functioning as a scaffold that organizes proton conductive keratan sulfate molecules.

By comparing the proton conductivity of native and proteinase K-digested ratfish gel, we learned that proteins may not play a direct role in the conductivity of the material, despite their clear contribution to the gel’s observed stiffness. In prior work (8), we speculated that sulfated polysaccharides, namely keratan sulfate molecules, were responsible for conferring the observed proton conductivity of AoL gel. If we consider the gel as a mixture of highly conductive polysaccharides (30) and less conductive proteins, removing the proteins should lead to a more conductive material as observed here. Nonetheless, the fact that the digested gel is more conductive than native gel confirms that proteins are not directly responsible for the observed proton conductivity. These data, along with the SAXS spectra, do not indicate which specific proteins play a role in the gel’s structure and function. However, our results suggest that proteins do confer the gelatinous nature of the material and contribute to the maintenance of the structured polymer network but are not critical for proton conductivity.

Finally, we note that our proposed model (Figure 3D) for the structure of the AoL gel is intrinsically different from that given by Zhang *et al*. (9). This is, in part, due to differences in the data types employed. The Zhang *et al*. model was generated by inference using proteomics data from the AoL gel and presumptive scaffolding of keratan sulfate, conceptually similar to the bottle-brush model of aggrecan (31). In contrast, our model is based on AFM and SEM imaging along with corroborating SAXS spectral analysis and proposes a colloidal gel organization. Although we never observed anything reminiscent of bottle-brush structures in AFM, it will be important to subject both models to further physicochemical and structural analyses in order to better deduce the underlying electrosensory mechanisms of the AoL of cartilaginous fishes.

## Materials & Methods

### Biological specimens

An adult little skate (Leucoraja erinacea) specimen was obtained from Marine Biological Laboratory and euthanasia was carried out using tricaine methylsulfonate (Western Chemical Inc., #NC0872873) at 1000 mg/L and conducted under the approved protocol: IACUC16-014 (Benaroya Research Institute). Expired and frozen adult spotted ratfish (*Hydrolagus colliei*) specimens were obtained from Moss Landing Marine Laboratories. The specimens were dethawed in a refrigerator overnight then AoL gel was removed by applying pressure to the pores with gloved fingers or the blunt side of a scalpel, placed in tubes, and stored at −80°C until use.

### CBD staining and fluorescence microscopy

AoL from an adult *L. erinacea* specimen were dissected from the hyoid cluster and labeled with CBD and DAPI following protocols described in (12). Whole-mount tissues were imaged using a Leica M205FA fluorescent stereoscope equipped with a DFC360FX monochrome CCD camera and a DFC425C color CCD camera.

Wax sections of formalin-fixed *H. colliei* tissues were prepared using standard processing, embedding and microtomy protocols and stained following protocols described in (12), and epifluorescent images were taken using either a Leica DMR upright epifluorescent microscope equipped with a SPOT RT Slider cooled 1.4 megapixel color/monochrome CCD camera and an Insight 4 megapixel color CCD camera (Diagnostic Instruments).

An aliquot of AoL gel from *H. colliei* was placed in a small centrifuge tube. Chitin-binding probes (CBD) (12) were diluted in 1X PBS and pipetted gently on top of the gel sample and incubated overnight at 4°C. The next day, the CBD liquid was gently replaced with Milli-Q water to wash. Gel was pulled from the tube, placed on a slide then imaged with a Leica DMP fluorescence microscope with a Qimaging Retiga Exi CCD camera.

### Gel digestion with proteinase K (for AFM imaging) & Analysis by SDS PAGE

Equal volumes of AoL gel and water were combined in a tube and vortexed vigorously. Proteinase K was added to a final concentration of 40 *μ*g/*μ*L and incubated for one hour at 50°C. An aliquot was then added to dialysis tubing with MWCO 12-14 KDa (Fisher #21-152-15), dialyzed overnight in 5 L of water at 4°C, then stored at −20°C until use. For SDS PAGE, 30 *μ*L of gel sample was combined with 10 *μ*L of reducing buffer and incubated at 95°C for 5 minutes. 1X running buffer was prepared and 1% SDS added. Samples were loaded into precast gels (Sigma #PCG2001-10EA) along with marker (Lonza ProSieve Color Protein Marker #BMA50550). Gel was run at 50 V for 20 minutes, then at 100 V for 50 minutes, then rinsed with DI water. Coomassie Brilliant Blue stain (Fisher #501035935) was poured over gel and incubated for 5 hours. Destain was then added to gel and swirled overnight before pictures were taken.

### Gel digestion with proteinase K (for SAXS)

Equal volumes of AoL gel and 1X PBS were combined in a tube and vortexed vigorously. Proteinase K was added to the gel sample to a final concentration of 0.1 *μ* g/*μ* L and incubated at 50°C for one hour. Sample was then dialyzed with 3.5 KDa dialysis tubing in 2 L of water for 5 hours.

### Gel digestion with proteinase K (for sonication & AFM)

Equal volumes of AoL gel and buffer (0.1 M Tris + 0.05 M EDTA + 0.1% SDS) were combined and proteinase K was added to a final concentration of 40 *μ*g/*μ*L. The solution was incubated at 50°C for an hour, then heated at 60°C for 30 min to denature remaining enzymes. The resulting solution was dialyzed overnight in 5 L of water with 12-14 KDa dialysis tubing (Fisher #21-152-15). Lastly, the sample was sonicated on the continuous setting on ice for 20 minutes.

### Ultracentrifugation of AoL gel

Several milliliters of AoL gel were dialyzed overnight in 3.5 KDa dialysis tubing in 5 L of water at 4°C to remove salts. A thick-walled polycarbonate lidless centrifuge tube was filled with dialyzed gel (∼500 *μ*L) then spun at ∼300,000-400,000 x g for 16 hours at 4°C with a TLA-120.1 Beckman Coulter rotor in an Optima Max-XP ultracentrifuge. A sample of Proteinase K-digested gel was put into a centrifuge tube and also spun at ∼300,000-400,000 x g for 16 hours.

### Synchrotron x-ray experiments

Small angle x-ray scattering (SAXS) experiments were carried out at Lawrence Berkeley National Laboratory’s (LBNL) Advanced Light Source (ALS) on beamline 7.3.3. Synchrotron radiation provides a high intensity, collimated x-ray beam, ideal for studying poorly ordered, dilute isotropic aqueous materials such as biological gels. Previously prepared gel samples were inserted into 1.5 mm quartz x-ray capillaries (Charles Supper Inc.) by gentle centrifugation and mounted in the beam path in a transmission configuration. Capillaries were mounted in a motorized translation stage, allowing precise control of capillary position in a plane perpendicular to the beam direction. A water filled capillary was included to obtain scattering data for the water background of the gels. A capillary containing silver behenate was included for beam center calibration and determining the sample to detector distance which was 3529.37 mm. To perform the scattering experiments, we used an approximately 300 *μ*m (H) x 700 *μ*m (W) beam at 10 keV and exposed the gels for 2.0 seconds per capillary at several different positions to obtain the most intense scattering pattern for analysis. The scattering patterns were recorded on a Pilatus 2M detector for analysis. The pixel size of the detector was 172 *μ*m^2^.

Data analysis was performed using the Nika and Irena macros in Igor Pro by WaveMetrics. The patterns showed diffuse scattering and rings of approximately uniform intensity (no significant alignment was observed) and were radially integrated to obtain 1D intensity plots as a function of q, the scattering vector. To perform the integration, we selected sectors on the scattering pattern image at various azimuthal angles and angular widths. The intensity plots shown were taken from the integration of sectors at an azimuthal angle of 60 degrees with an angular width of 20 to 40 degrees. The Igor wave arithmetic tool was used to subtract the water background.

### Supercritical drying

An AoL gel sample was placed in 3.5 KDa dialysis tubing, sealed on both ends using dialysis clamps, and left in 200 proof ethanol for several hours (in most cases, overnight). Dialysis tubing became very stiff and little liquid remained inside after ethanol incubation. Often, white precipitate was observed coating the inside of the tube. Dialysis clamps were replaced with twist-ties before introduction to the supercritical drying system. Supercritical drying was performed with a Denton Vacuum, Inc supercritical drying system. The chamber was flushed with liquid CO_2_ until no ethanol smell was observed in the effluent followed by a 2-hour soak in pressurized liquid CO_2_ to allow exchange of ethanol inside the dialysis tubing. Phase change to supercritical CO_2_ was achieved by heating the chamber to ∼50°C (reaching pressure of ∼1400 psi). Final drying was achieved by reducing sample chamber pressure allowing the supercritical CO_2_ to transition to the gas phase. Samples were stored under vacuum until imaging.

### Scanning Electron Microscopy (SEM)

AoL gel samples were ambiently dried on a freshly cleaved piece of mica attached to an aluminum stub using double-sided carbon tape. For supercritically dried samples, a piece of double-sided carbon tape was stuck onto aluminum stubs then pressed onto the inside of the dialysis tubing containing supercritically dried samples. Scanning electron microscopy was performed with the Zeiss Gemini500 FEG-SEM at the Imaging and Microscopy Facility of University of California, Merced. Beam landing energy of 500 eV and secondary electron detection were used.

### Atomic Force Microscopy (AFM)

AoL samples were dropped directly onto freshly cleaved sheets of mica and dried ambiently in a covered box for several hours. AFM was performed using a Veeco Innova instrument in tapping mode. Probes with a spring constant of 40 N/m (Tap300Al-G, Budget Sensors) were used to image air-dried samples drop-cast on freshly cleaved mica to obtain topographical, amplitude and phase images in air. Images were collected at a scan rate of 1 line per second at room temperature.

### Proton conductivity

Samples of dialyzed gel and proteinase-K treated dialyzed gel were brought to UC Santa Cruz where analyses were performed. The two-terminal devices used in EIS measurements were fabricated on a glass wafer (Figure 4A). Prior to device fabrication, the substrates were cleaned by sequential sonication in acetone, IPA and water. Then, a 10 nm Titanium adhesion layer overlaid with a 100 nm gold was electron-beam evaporated onto the clean substrates through a shadow mask. The dimensions of the paired electrodes were 1 cm wide by 2 cm long with an inter-electrode separation of 50 µm. A polydimethylsiloxane (PDMS) well was made with a 4 mm biopsy punch and bonded to the glass wafer to define the geometry of the gel. The devices were completed by drop casting the native *H*.*colliei* AoL gel and digested gel directly into the PDMS well, and the resulting films were allowed to dry in air (Supplemental Figure 5). We hydrated the samples by incubating them in a home-made sealed chamber at 90% relative humidity (RH) at room temperature for 2 hours with D_2_O or H_2_O. After incubation, Nyquist plots were recorded by Autolab PGSTAT128N between 100 kHz and 0.1 Hz at 10 mV amplitude and analyzed by Nova 2.0 software. Then, we calculated the conductivity (*σ*) using the following equation: σ = *L* / *R*_b_ *A* where *A* is the cross-sectional area given by the width of the contact (4 mm) multiplied by the thickness of the sample as measured with atomic force microscopy (2.7 ± 1.9 *μ*m for ratfish gel, 0.46 ± 0.27 *μ*m for digested ratfish gel), *L* is the device length or electrode separation (50 *μ*m), and *R*_b_ is the value of resistance obtained from the equivalent circuit we mentioned above.

## Acknowledgments

Thanks to Jason Cope (NOAA), Scott Hamilton, & Matthew Jew (Moss Landing Marine Laboratories) for providing access to expired *H. colliei* specimens for AoL gel extraction. This research used beamline 7.3.3 of the Advanced Light Source, which is a DOE Office of Science User Facility under contract no. DE-AC02-05CH11231 (32). Funds were also provided by UC Merced (to CTA) and by a UC Merced Senate award to CTA and LSH.

## Notes

**Competing Interest Statement:** The authors declare no competing interests.

### Competing Interest Statement

The authors have declared no competing interest.

## References

1. Kalmijn AJ (1974) The Detection of Electric Fields from Inanimate and Animate Sources Other Than Electric Organs. Handbook of Sensory Physiology: Electroreceptors and Other Specialized Receptors in Lower Vertebrates), Vol 3, pp 147–200.

2. Tricas TC, Michael SW, & Sisneros JA (1995) Electrosensory optimization to conspecific phasic signals for mating. Neuroscience Letters 202:129–132.

3. Newton KC, Gill AB, & Kajiura SM (2019) Electroreception in marine fishes: chondrichthyans. J Fish Biol 95(1):135–154.

4. Kempster RM, McCarthy ID, & Collin SP (2012) Phylogenetic and ecological factors influencing the number and distribution of electroreceptors in elasmobranchs. J Fish Biol 80(5):2055–2088.

5. Raschi W (1986) A Morphological Analysis of the Ampullae of Lorenzini in Selected Skates (Pisces, Rajoidei). Journal of Morphology 189:225–247.

6. Murray RW & Potts WTW (1961) The Composition of the Endolymph, Perilymph, and Other Body Fluids of Elasmobranchs. Comparative Biochemistry and Physiology 2:65–75.

7. Doyle J (1967) The ‘Lorenzan Sulphates’ - A New Group of Vertebrate Mucopolysaccharides. Biochemical Journal 103:325–330.

8. Josberger EE, et al. (2016) Proton Conductivity in ampullae of Lorenzini jelly. Science Advances 2:e1600112.5):1-6.

9. Zhang X, et al. (2018) Structural and Functional Components of the Skate Sensory Organ Ampullae of Lorenzini. ACS Chem Biol.

10. How MJ & Jones JVS (1969) Comparative Studies of Lorenzini Jelly from 2 Species of Elasmobranch .I. Preparation of Glycopeptides. Carbohydrate Research 11(2):207-&.

11. How MJ, Jones JVS, & Stacey M (1970) Comparative Studies of Lorenzini Jelly From Two Species of Elasmobranch. Carbohydrate Research 12:171–181.

12. Phillips M, et al. (2020) Evidence of chitin in the ampullae of Lorenzini of chondrichthyan fishes. Current Biology 30(20):R1254–R1255.

13. Djabourov M (1988) Architecture of gelatin gels. Contemporary Physics 29(3):273–297.

14. Tang WJ, Fernandez J, Sohn JJ, & Amemiya CT (2015) Chitin is endogenously produced in vertebrates. Curr Biol 25(7):897–900.

15. Hirst LS, Pynn R, Bruinsma RF, & Safinya CR (2005) Hierarchical self-assembly of actin bundle networks: gels with surface protein skin layers. J Chem Phys 123(10):104902.

16. Buchtova N & Budtova T (2016) Cellulose aero-, cryo- and xerogels: towards understanding of morphology control. Cellulose 23:2585–2595.

17. Ganesan K, Dennstedt A, Barowski A, & Ratke L (2016) Design of aerogels, cryogels and xerogels of cellulose with hierarchical porous structures. Materials and Design 92:345–355.

18. Ganesan K, et al. (2018) Review on the Production of Polysaccharide Aerogel Particles. Materials (Basel) 11(11).

19. Quignard F, Valentin R, & Di Renzo F (2008) Aerogel materials from marine polysaccharides. New Journal of Chemistry 32(8):1300–1310.

20. Sahin I, Ozbakir Y, Inonu Z, Ulker Z, & Erkey C (2017) Kinetics of Supercritical Drying of Gels. Gels 4(1).

21. Stribeck N (2007) X-Ray Scattering of Soft Matter (Springer-Verlag, Berlin) 1 Ed p 240.

22. Rubinstein M & Colby RH (2003) Polymer Physics (Oxford University Press, Oxford).

23. Phillips R (2018) Membranes by the Numbers. Physics of Biological Membranes, eds Bassereau P & Sens P (Springer Nature, Switzerland), pp 73–105.

24. Bureekaew S, et al. (2011) One-dimensional imidazole aggregate in aluminium porous coordination polymers with high proton conductivity. Materials For Sustainable Energy: A Collection of Peer-Reviewed Research and Review Articles from Nature Publishing Group, (World Scientific), pp 232–237.

25. Soboleva T, et al. (2008) Investigation of the through-plane impedance technique for evaluation of anisotropy of proton conducting polymer membranes. Journal of Electroanalytical Chemistry 622(2):145–152.

26. Guo Q-H, et al. (2020) Single-Crystal Polycationic Polymers Obtained by Single-Crystal-to-Single-Crystal Photopolymerization. Journal of the American Chemical Society.

27. Jalili J & Tricoli V (2014) Proton conductance at elevated temperature: formulation and investigation of poly (4-styrenesulfonic acid)/4-aminobenzylamine/phosphoric acid membranes. Frontiers in Energy Research 2:28.

28. Haas J, Ravi Kumar MN, Borchard G, Bakowsky U, & Lehr CM (2005) Preparation and characterization of chitosan and trimethyl-chitosan-modified poly-(epsilon-caprolactone) nanoparticles as DNA carriers. AAPS PharmSciTech 6(1):E22–30.

29. Zeng JB, He YS, Li SL, & Wang YZ (2012) Chitin whiskers: an overview. Biomacromolecules 13(1):1–11.

30. Selberg J, Jia M, & Rolandi M (2019) Proton conductivity of glycosaminoglycans. PLoS One 14(3):e0202713.

31. Harder A, Walhorn V, Dierks T, Fernandez-Busquets X, & Anselmetti D (2010) Single-molecule force spectroscopy of cartilage aggrecan self-adhesion. Biophys J 99(10):3498-3504.

32. Hexemer A, et al. (2010) A SAXS/WAXS/GISAXS Beamline with Multilayer Mnochromator. Journal of Physics: Conference Series 247.

