## Supplementary Information for "Structural Characteristics and Proton Conductivity of the Gel Within the Electrosensory Organs of Cartilaginous Fishes"

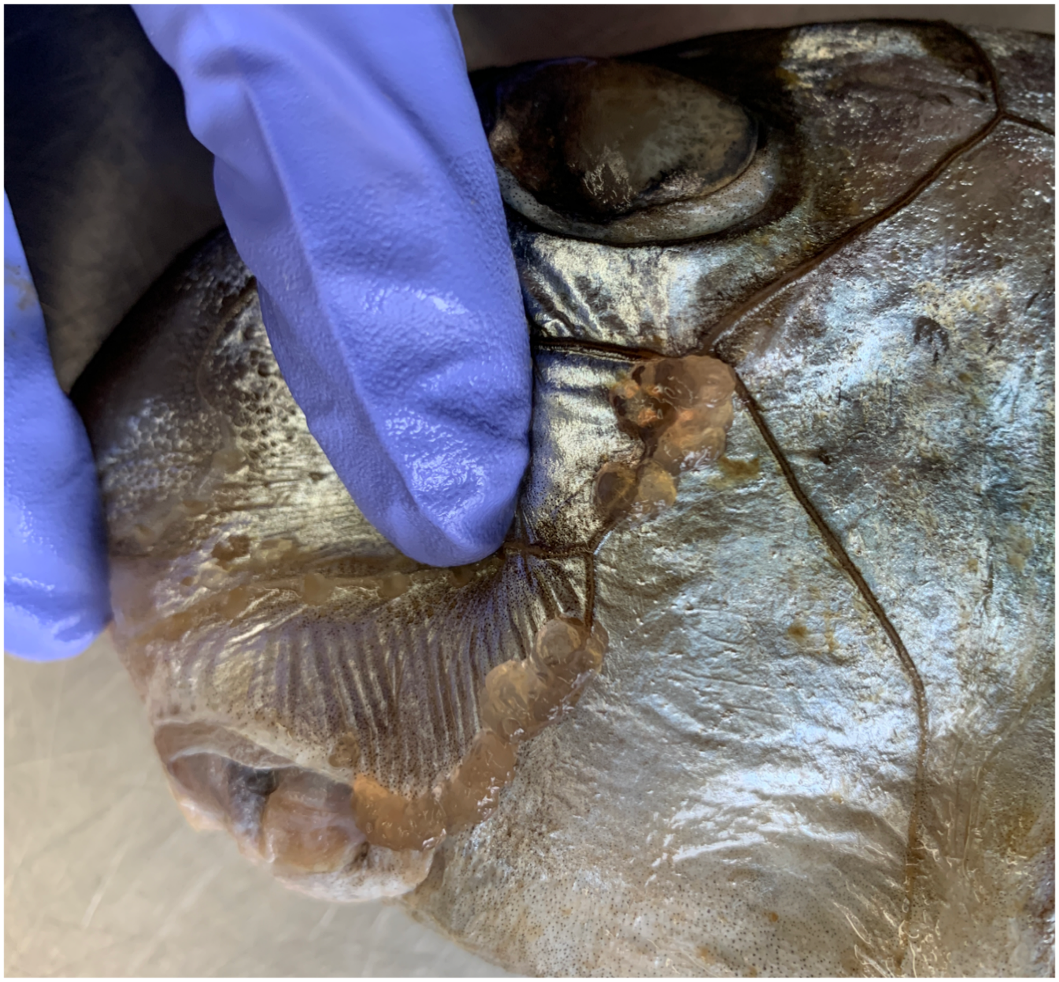

**Supplemental Figure 1** AoL gel removal process. Image that shows hydrogel being squeezed from the AoL pores of an expired spotted ratfish (*Hydrolagus colliei*) specimen. Thumb (in purple glove) applies pressure to the canals between the eye (top) and mouth (bottom). Gel is pink in color due to blood and cellular contaminants resulting from the freeze-thaw process and mechanical expulsion.

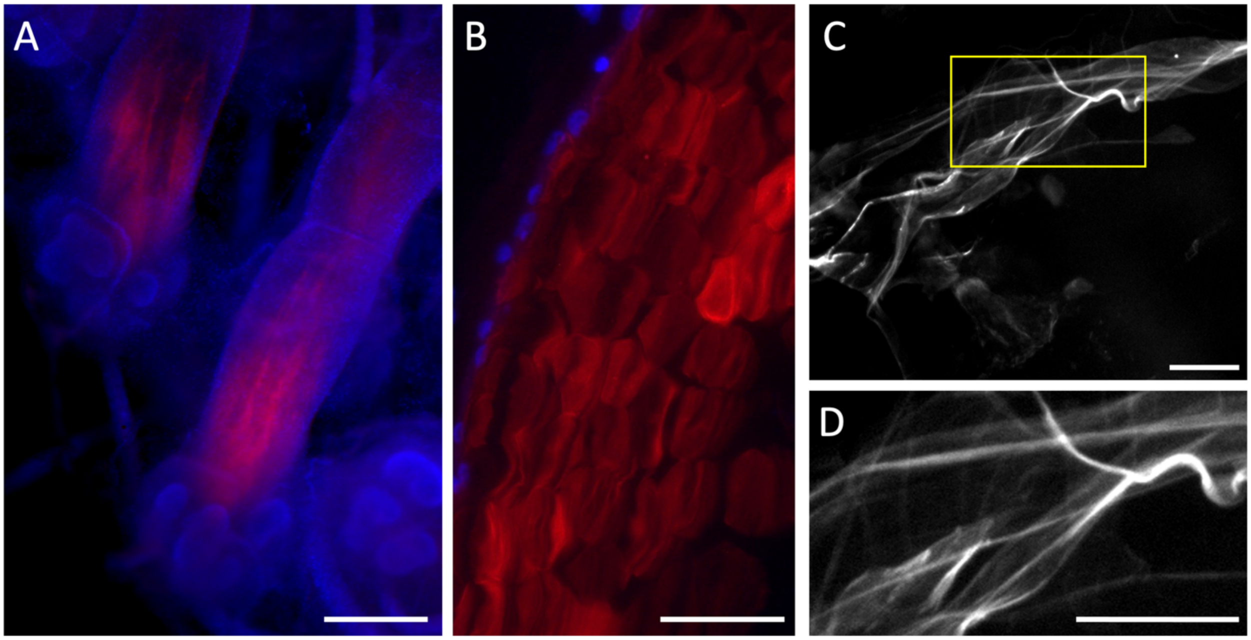


**Supplemental Figure 2** Visualization of AoL gel via fluorescence microscopy. **A)** Formalin-fixed whole-mount tissues from little skate (*Leucoraja erinacea*) specimens imaged with a fluorescence stereomicroscope. Tissues were stained with chitin-binding CBD probes (red) as well as nuclear stain (DAPI, blue). The outlines of two AoL are revealed by DAPI and CBD is shown to label the gel within the tubular organs. See Figure 1B for the overall anatomy of AoL. **B)** Formalin-fixed sectioned tissue from spotted ratfish (*H. colliei*) labeled with CBD (red) and DAPI (blue) and imaged with fluorescence microscopy. Image shows a cross section through an AoL with DAPI-labeled nuclei lining the edge of the canal (upper left) and CBD-labeled gel filling the acellular canal lumen. **C, D)** Aqueous unfixed AoL gel from *H.colliei* labeled with CBD (white) and imaged with fluorescence microscopy. Area delineated by the yellow rectangle in (C) is shown at higher magnification in (D). Scale bars – A: 500 $\mu$m; B-D: 50 $\mu$m.

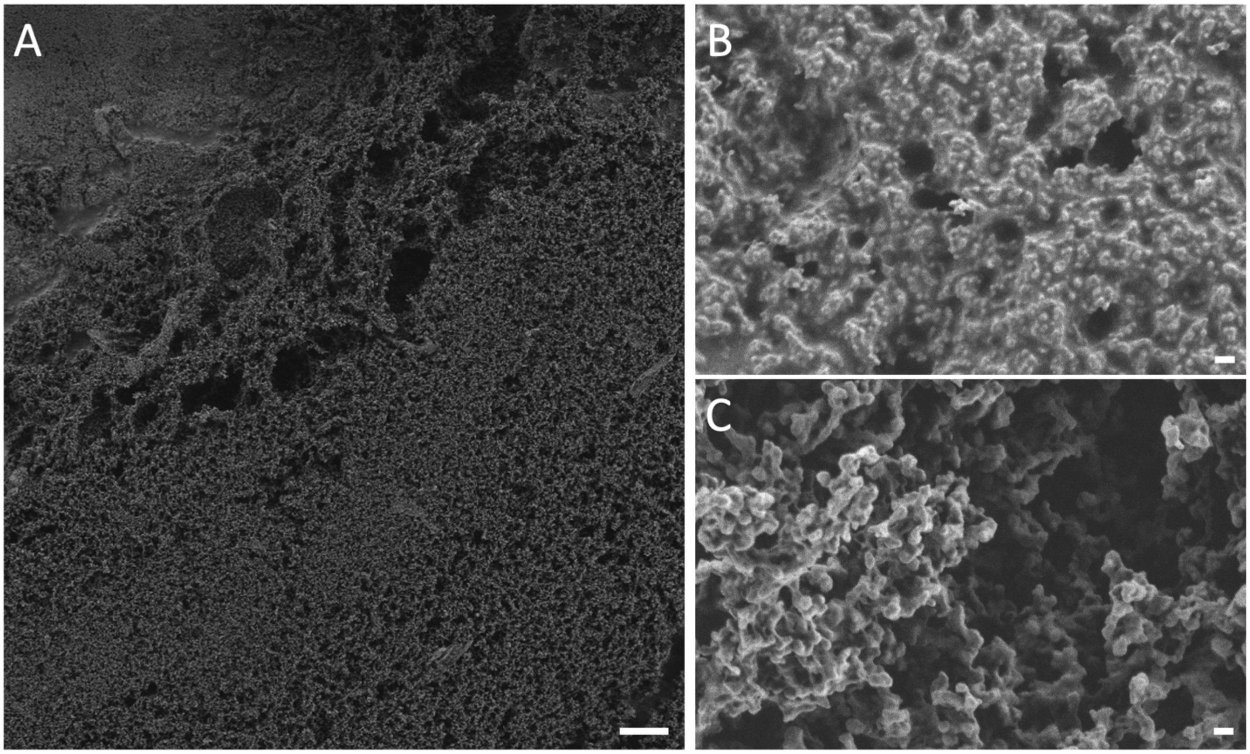


**Supplemental Figure 3** SEM images of supercritically dried *H. colliei* AoL gel. **A)** Low magnification demonstration of the texture of supercritically dried AoL gel. **B, C)** High magnification images of AoL gel in two different areas of the sample. There was some variability in the texture and order of the structures. In a few areas, the structure appeared dense and tightly packed (B), but the majority of the material demonstrated marked porosity (C), as is typical of aerogels (1). Scale bars – A: 10 $\mu$m; B, C: 200 nm.

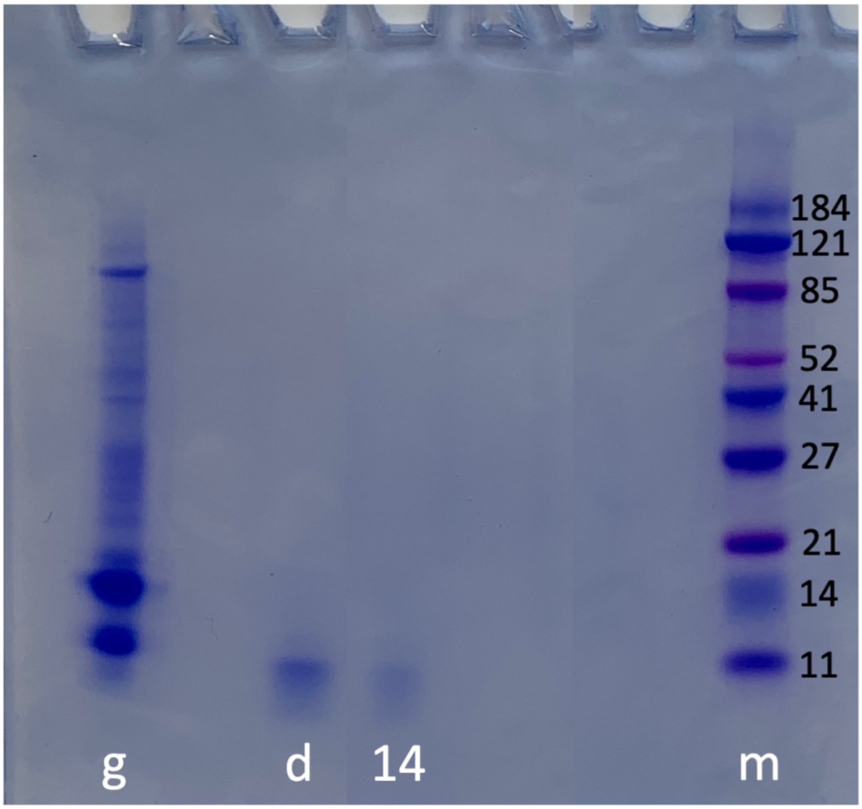


**Supplemental Figure 4** Image of AoL gel samples electrophoresed through an SDS polyacrylamide gel (PAG) and stained with Coomassie Blue. The lanes contain native *H. colliei* AoL gel (g), gel digested with proteinase K (d), digested gel that had been dialyzed overnight in 12-14 KDa dialysis tubing (14), and a marker (m). Molecular weights in KDa are shown in black next to the marker. These data show that AoL gel contains proteins of many sizes and that proteolytic digestion is effective at breaking the large proteins into smaller peptides that are <11 KDa. Dialysis is effective at further removing residual peptides.

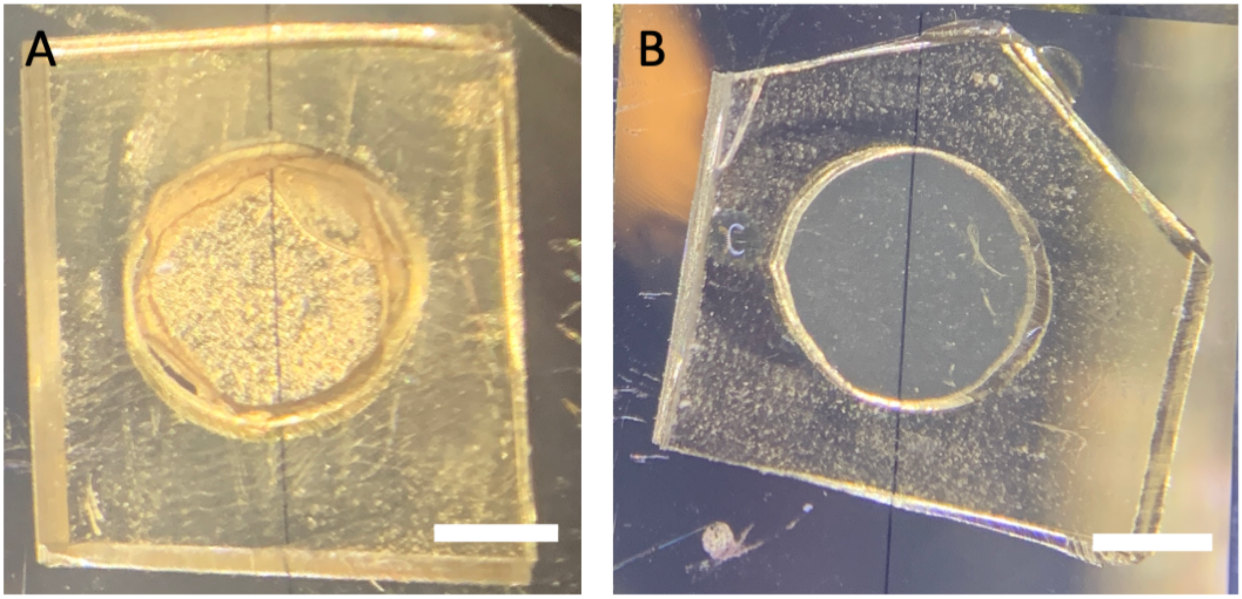


**Supplemental Figure 5** *H. colliei* AoL gel samples dried on proton conductivity devices inside a circular spacer. **A)** Optical image of the native gel on the device. **B)** Optical image of the digested gel on the device. Scale bars – 2 mm.

### SI References

1. Ganesan K, Dennstedt A, Barowski A, & Ratke L (2016) Design of aerogels, cryogels and xerogels of cellulose with hierarchical porous structures. *Materials and Design* 92:345-355.
